# Resilience not yet apparent in soil fungal communities of the boreal forest from one to five years after wildfire across a severity gradient

**DOI:** 10.1101/2025.03.29.646032

**Authors:** Thea Whitman, Jamie Woolet, Miranda C. Sikora, Dana B. Johnson, Denyse A. Dawe, Ellen Whitman

## Abstract

Wildfires are natural disturbances characteristic of boreal forests. However, fire regimes around the world are changing, with effects on fire frequency and fire severity. Here, we present data from one and five years post-fire across 40 different sites from the boreal forest of the Northwest Territories and Alberta, Canada, asking the following questions: (1) Do the factors that structure post-fire soil fungal communities change over time? (2) Is there evidence for resilience in fungal community composition across different burn severities? (3) Do fire-enriched taxa change one vs. five years post-fire? Factors correlated with fungal community composition remain largely the same five years post-fire, with a declining correlation with burned/unburned, and an increasing association with vegetation community composition. This suggests that the immediate effects of fire on fungal community composition wane or diverge with time, while the influence of longer-term effects of fire, such as changes in vegetation, increases. Fungal communities failed to demonstrate resilience in community composition five years post-fire, in contrast to our findings for bacterial communities, and suggesting that fungal communities may be more closely tied to soil properties and vegetation communities that take longer to recover post-fire. Finally, we identify and classify fire-responsive fungi into different response types, based on their enrichment or depletion one vs. five years post fire. Persistent fire responders include taxa from the genera *Penicillium*, *Coniochaeta*, and *Calyptrozyma*, and the family *Venturiaceae*, but different taxa within a single genus respond differently to fires, underscoring that generalizations even at relatively fine taxonomic levels may be inappropriate, and that the mere presence of traits that may be relevant to post-fire success are insufficient alone to guarantee post-fire abundance. Future manipulative and observational studies will help us continue to dissect the multiple factors and traits structuring fungal responses to fires.

## 1. Introduction

Wildfires represent a fundamental natural disturbance in boreal forests. However, as a result of climate change, fire regimes in many parts of the world are changing, with increasing fire activity in response to more severe fire weather (Hanes et al., 2019; Jain et al., 2022), and increasing fire severity (Parks and Abatzoglou, 2020; E. Whitman et al., 2022). Many organisms have developed adaptations that allow them to thrive in fire-affected ecosystems, such as the serotinous cones of jack pine (Radeloff et al., 2004) or *Melanophila* beetles, which lay their eggs in fire-killed wood (Schmitz and Bousack, 2012). While fire ecology is relatively well-elucidated for above-ground organisms, our understanding of the effects of fires on soil organisms such as soil fauna, bacteria, and fungi, continues to emerge (Zaitsev et al., 2016). Understanding how soil microbes respond to wildfires across a range of conditions is of interest, to better understand the fundamental ecology of these organisms, and how changes in their communities may affect or reflect the state and functioning of the broader ecosystem.

Fungal fire ecology offers an interesting contrast to bacterial fire ecology, in that specific fire-associated fungi have been known for centuries (Seaver, 1909). Known by various names — fireplace fungi, phoenicoid, pyrophilous, anthracophilous, carbonicolus, or post-fire fungi (Fox et al., 2022) — certain fungi exhibit traits that enable their proliferation after fires. Historically, understanding of these fungi relied on observable sporocarps, but molecular techniques have revealed greater soil fungal diversity and raised questions about pyrophilous fungi ecology: What drives a taxon to fruit, proliferate, or dominate post-fire communities? Reflecting their diversity, fungi employ multiple strategies, categorized by Fox et al. (2022) as “fire-adapted,” “fire-responsive,” and “fire-resistant.” These strategies operate within the dynamic post-fire environment, shaped by ecosystem and burn characteristics over time.

Immediately following a fire, widespread belowground mortality—including fungi, bacteria, archaea, soil fauna, and plants—opens ecological space and provides resources such as lysed microbial cells and dead plant material for saprotrophs. Fire also combusts and transforms organic matter, depleting certain resources and altering the chemical and physical soil environment (Certini, 2005). Shifts in ecological interactions emerge as some organisms perish while others survive or recolonize, changing competition dynamics and symbiotic relationships. These diverse fire effects highlight the importance of varied fungal strategies across ecosystems and timescales. Together, these strategies determine whether fungal communities exhibit resilience after wildfire. Resilience can be described as the ability of a system to return to pre-disturbance states after a perturbation (Holling et al., 1973), and informs us about possible future states, including how the system may respond to changing disturbance and climate regimes.

Despite the wide-ranging potential fungal responses to fire in boreal forests, a few trends emerge. First, fungal biomass is reduced after wildfires, often more than that of bacteria (Pressler et al., 2018). Second, a number of studies have noted decreases in the relative abundance of ectomycorrhizal fungi after a wildfire, likely due to the death of host trees (Hewitt et al., 2022). At the same time, some studies have found increases in saprotrophs after wildfires (Fox et al., 2022), presumably due to the pulse of substrates. Broadly, these trends are mirrored by a decrease in fungi from the Basidiomycetes phylum, and an increase in those from the Ascomycetes phylum (Cairney and Bastias, 2007; Holden et al., 2016; Nelson et al., 2022). Molecular techniques have also allowed for the identification of specific taxa that are enriched post-fire. These include both macroscopically observed taxa, such as species from the genera *Pyronema* or *Morchella*, as well as those that are more commonly detected via molecular assays, such as species from the genus *Penicillium*.

Despite our improving understanding of fungal fire ecology, much of what we currently know is limited to either repeatedly sampled but short-term (1-2 year) (Pulido-Chavez et al., 2021; Ammitzboll et al., 2022; Pulido-Chavez et al., 2023) or single-timepoint studies. Our understanding of how communities change over time is largely based on chronosequence studies, which offer valuable information, but may conflate spatial variation with that due to time since fire. Studies that tracked fungal communities over years to decades include Salo et al. (Salo and Kouki, 2018; Salo et al., 2019), who tracked sporocarps for 12-years post-fire in Finland, and Franco-Manchón et al. (Franco-Manchón et al., 2019), who observed sporocarps 1 and 5 years post-fire in Mediterranean and boreal pine forests. Franco-Manchón et al. (2019) found no mycorrhizal fungi one year post-fire, and only a few taxa five years post-fire. In contrast, Salo and Kuki (2018) observed ectomycorrhizal fungi increasing following moderate severity burns, but declining following high-severity burns, highlighting the importance of considering fire in the context of its severity, not as a binary treatment. Taylor et al. (2010) offered an early perspective on post-fire fungal recovery, although this paper focused primarily on seasonal and yearly community compositional shifts, and on sites that were already at least 23 years post-fire. Applying a modern molecular approach to investigate longer-term fungal response across a gradient of burn severity remains a critical research need.

In 2015, we assessed the effects of fire on soil fungal communities one year post-fire across a burn severity gradient in the boreal forest of northern Canada(T. Whitman et al., 2019). In the present study, we add a 5-year timepoint to our previous 1-year post-fire study, allowing us to investigate how fungal communities change over intermediate timescales post-fire by tracking the same sites over time, rather than using a chronosequence approach. We sought to ask a series of questions about how the community changes after a burn:

1. Do the factors that structure post-fire soil fungal communities change between one and five years? We predicted that the same factors would structure fungal community composition both years, with a waning influence of whether the site was burned.
2. Is there evidence for resilience in fungal community composition, and does it differ with different burn severities? We predicted that there would be a similar pattern to that observed in the bacterial/archaeal dataset, where burned communities would become more similar to unburned communities five years post-fire, and that this would not vary with burn severity.
3. Do the taxa that are enriched post-fire change between one and five years? We predicted that the strongest responders (including *Penicillium*, *Calyptrozyma*, *Coniochaeta*, and *Fusicladium*) would remain significantly enriched five years post-fire. We also sought to classify fire-responsive taxa into different response groups, based on their different responses one and five years post-fire.

## 2. Methods

### 2.1 Study region and site selection

Our study region is in the southern half of the boreal and taiga plains ecoregions of northwestern Canada (northern Alberta and the southern Northwest Territories). The study region has long, cold winters and short, warm summers, with mean annual temperatures between −4.3 °C and −1.8 °C and annual precipitation ranging from 300 to 360 mm (ESWG, 1995; Wang et al., 2012). The region’s fire regime is characterized by infrequent stand-replacing fires every 40-350 years on average (Boulanger et al., 2012) and, due to its small and dispersed human population, fires are often managed with minimal suppression and control, when appropriate. 2014 was an exceptional year for wildfires in the region (Whitman et al., 2018b), and offered an opportunity to study wildfires across a range of vegetation communities and burn severities. We selected 40 sites in the Northwest Territories and northern Alberta (Wood Buffalo National Park), Canada, which spanned low to high burn severities and included unburned sites. We sampled them one year post-fire, in 2015, and again five years post-fire, in 2019 (Figure 1). The fires and the drivers of burn severity are described in detail in Whitman et al. (2018b), their effects on understory vegetation are described in detail in Whitman et al. (2018a), their one-year effects on soil microbial communities are described in detail in T. Whitman et al. (2019), and their five-year effects on soil bacterial communities are described in detail in (T. Whitman et al., 2022). This paper compares fungal community response one year vs. five years post-fire and is a companion to the Whitman et al. (2022) bacterial/archaeal dataset. The two datasets were not prepared and published together due to delays in fungal sequencing and analysis related to the global COVID-19 pandemic.

**Figure 1.**
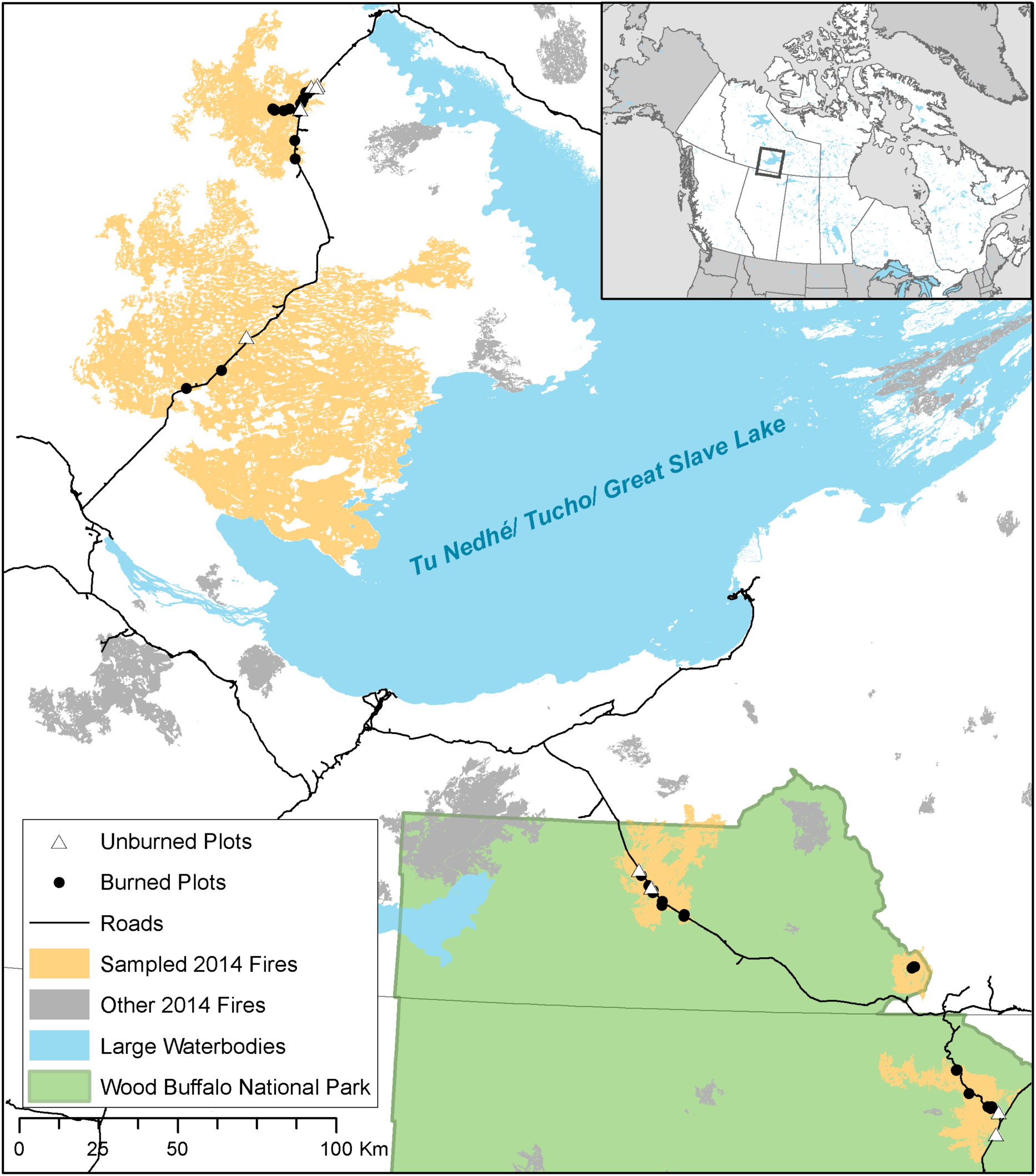
Study region of northern Alberta and the Northwest Territories, Canada, including Wood Buffalo National Park (WBNP – green shading). Closed circles indicate burned plots, open triangles indicate unburned plots, yellow shapes indicate the total extent of sampled fires, and grey shapes indicate other 2014 fires in the region. Inset indicates relative location within Canada.

The six wildfires in this study were very large (14,000 to 700,000 ha). The soils in these regions are mostly classified as Typic Mesisols (21 sites), Orthic Gleysols (6 sites), or Orthic Gray Luvisols (10 sites) (Soil Landscapes of Canada map v.3.2). They span a wide range of soil properties, with pH values ranging from 3.2 (treed wetlands) to 8.1 (uplands with calcareous soil), total C ranging from 0.5% (mineral horizon) to 52% (organic horizon), and a wide range of soil textures. Vegetation communities were classified as jack pine-dominated (*Pinus banskiana* Lamb.) uplands, black spruce-dominated (*Picea mariana* (Mill.)) uplands, mixedwood uplands with both coniferous and broadleaf trees, or treed wetlands (Beckingham and Archibald, 1996).

### 2.2 Site assessment methodologies

Sites were selected and characterized as described in detail by Whitman et al. (2018a; 2018b). Briefly, field sites were selected to represent the local range of burn severity and vegetation communities, resulting in a total of 31 burned field sites across three vegetation communities (pine, spruce, and mixedwood). An additional 9 control sites (not burned within the last 38 years before sampling, mean time since fire 95 years; “unburned”) were selected, chosen to reflect the range of vegetation communities sampled in the burned plots, for a total of 40 sites. These represent 40 of the 62 original sites (we excluded non-forested sites from revisitation, and some sites were not accessible five years post-fire or destroyed by human activity). At each site, one year post-fire, we established a 30 × 30 m square plot with 10 × 10 m subplots at the four corners. We assessed burn severity in the four subplots using burn severity index (BSI (Loboda et al., 2013); described in detail in Whitman et al. (2018a)). We returned to sites five years post-fire to re-assess vegetation composition (reported in (Dawe et al., 2022)) and sample soils.

At each plot, we took soil cores (5.5 cm diameter, 13.5 cm depth) at three locations. The sampling scheme was modified between sampling years in order to optimize efficient representation of site characteristics, including vegetation: at one year post-fire, samples were taken at plot centre, 7 m SW of centre, and 7 m NE of centre, and at five years post-fire, samples were taken at plot centre, 17.5 m N of centre, and 17.5 m S of centre (these changes are not expected to affect our central findings, as indicated by comparing unburned sites between years). Soil cores were gently extruded and separated into organic (O) horizons (where present) and mineral (M) horizons (where present in the top 13.5 cm of soil profile). Where mineral horizons were present, one year post-fire, they were sampled to whatever depth represented the bottom of the 13.5 cm core.

Five years post-fire, we modified the protocol so a consistent depth (5 cm) was sampled for mineral soils. Although the two years’ sampling approaches would have rarely or never included different genetic soil horizons, this would undoubtedly affect certain soil properties that vary with depth. However, we do not expect that this methodological change drives our key findings, as indicated by broad similarities in data between sampling years and similar patterns observed in the O horizon samples, for which the protocol remained consistent between years.

The three samples were combined by horizon at each site and mixed gently by hand in a bag. From these site-level samples, sub-samples were collected for microbial community analysis and stored in LifeGuard Soil Preservation solution (QIAGEN, Germantown, MD) in a 5 mL tube (Eppendorf, Hamburg, Germany). Tubes were kept as cold as possible while in the field (usually for less than 8 h, but up to 2 days for remote sites) and then stored frozen. We do not expect these decisions to have large impacts on community composition (Lauber et al., 2010). The remaining soil samples were air-dried and analyzed for pH (1:2 soil:DI water for mineral samples; 1:5 soil:DI water for organic samples) and total C (combustion analysis). Soil texture was only measured one year post-fire (Whitman et al., 2019).

### 2.3 DNA extraction, amplification, and sequencing

DNA extractions were performed for each sample (71 soil samples total), with two blank extractions for every 24 samples (identical methods but using empty tubes, half of which were sequenced), using a DNeasy PowerLyzer PowerSoil DNA extraction kit (QIAGEN, Germantown, MD) following manufacturer’s instructions. (For the one year post-fire samples, duplicate DNA extractions were performed and sequenced. Duplicates were highly similar, so single extractions were performed for the five years post-fire samples.) Extracted DNA was amplified in triplicate PCR, targeting the ITS2 region with 5.8S-Fun and ITS4-Fun primers (Taylor et al., 2016) with barcodes and Illumina sequencing adapters added as per (Kozich et al., 2013) (all primers in Supplemental Table S1). The PCR amplicon triplicates were pooled, purified and normalized using a SequalPrep Normalization Plate (96) Kit (ThermoFisher Scientific, Waltham, MA). Samples, including blanks, were pooled and library cleanup was performed using a Wizard SV Gel and PCR Clean-Up System A9282 (Promega, Madison, WI). The pooled library was submitted to the UW Madison Biotechnology Center (UW-Madison, WI) for 2×300 paired end (PE) Illumina MiSeq sequencing.

### 2.4 Sequence data processing and taxonomic assignments

Sequencing data from both sampling years were prepared identically, and were re-processed for this study, following identical protocols, but working with each year separately due to potential run-to-run variation in error rates or other differences between years. We first trimmed all reads to ensure no primers were retained on short sequences using cutadapt (Martin, 2011) as implemented in QIIME2 (2023.5; Bolyen et al., 2019). We quality-filtered and trimmed, dereplicated, learned errors, determined operational taxonomic unites (OTUs), and removed chimeras from ITSxpress-trimmed ITS2 reads using dada2 (Callahan et al., 2016) as implemented in QIIME2 (2023.2; Bolyen et al., 2019). Taxonomy was assigned to the ITS2 reads using a QIIME2 (Bolyen et al., 2019) scikit-learn feature classifier (Bokulich et al., 2018) trained on the dynamic similarity cutoff UNITE fungal ITS database (Pruesse et al., 2007; Quast et al., 2013; Yilmaz et al., 2013; Abarenkov et al., 2022).

We then wanted to merge the two datasets to create a final, merged OTU table. Although the OTU clustering algorithm can distinguish taxa that differ by a single nucleotide and we used identical sequencing and bioinformatics protocols between the two years, we were concerned that merging only OTUs with 100% identical sequences and sequence lengths between the two years might be overly conservative. Fungi can have multiple copies of the ribosomal RNA operon in their genomes (for which there could be small variations in the ITS2 region), and isolates from the same recognized species can have small variations in their ITS2 region as well. Thus, we took the approach of first merging any OTUs with sequences that were 100% identical, and then, given the relatively high fraction of taxa with species-level taxonomic assignments (66% and 62% of OTUs in one-year and five-year datasets, respectively), we further merged OTUs that shared the same species name. We then removed all taxa for which we were unable to assign a phylum during the classification step, to eliminate any spurious eukaryotes. Due to relatively stringent quality trimming, the maximum OTU sequence length was 416 bp. We recognize that this means that we excluded any OTUs from the dataset that had longer amplicons, since we would expect this primer pair to include taxa with up to 511 bp length amplicons (Taylor et al., 2016). However, less stringent trimming, while allowing longer OTUs to be retained, resulted in substantially fewer reads overall being retained, due to low quality scores, and inconsistency between the two years’ datasets due to differences in quality score profiles. Our read processing approaches ultimately retained a mean of 70% and 75% of initial reads, for one and five years post-fire, respectively.

We classified all taxa by trophic modes using FunGuild (Nguyen et al., 2016) with the assigned UNITE taxonomy (Pruesse et al., 2007; Quast et al., 2013; Yilmaz et al., 2013), interpreting only classifications that were rated as being “highly probable”.

Throughout this paper, all analyses are performed with only the sites that were included in both years of sampling, so minor discrepancies (but no qualitative differences) are to be expected between the one-year data presented here and in the original paper (T. Whitman et al., 2019). In particular, we did not sample open wetlands five years post-fire, so these are not included in the dataset.

### 2.6 Statistical analyses

All analyses and plotting were done with R (Team, 2022) in RStudio, using packages *phyloseq* (McMurdie and Holmes, 2013), *dplyr* (Wickham et al., 2021), and *ggplot2* (Wickham, 2016).

To determine which site and soil sample parameters remained significant predictors of community composition five years post-fire, we ran a permutational multivariate analysis of variance (PERMANOVA) on Bray-Curtis dissimilarities (Bray and Curtis, 1957) with Hellinger-transformed relative abundances using the *vegan* package in R (Oksanen et al., 2021), reporting R^2^ and p-values for each year’s dataset. After considering our results, we wondered whether vegetation transitions might be associated with fungal community composition. To explore this idea, we ran a PERMANOVA comparing the predictive power (R^2^) of dominant overstory vegetation pre-fire and five years post-fire for fungal community composition, for upland burned sites where vegetation transitions had occurred.

To determine whether burned communities were more similar to unburned communities five years post-fire *vs*. one year post-fire, we compared Bray-Curtis dissimilarities on Hellinger-transformed relative abundances between the two years using a Mann-Whitney U test (because dissimilarities one year post-fire were not normally distributed). We plotted the dissimilarities using non-metric multidimensional scaling (NMDS). To determine whether community composition of severely burned sites changed more between one and five years post-fire, we tested whether Bray-Curtis dissimilarities on Hellinger-transformed relative abundances for paired one-year and five-year samples from the same site were significantly correlated with burn severity index using a Kruksal-Wallis test.

In order to identify which OTUs were significantly enriched (“positive response”) or depleted (“negative response”) in burned plots *vs.* unburned plots for each year individually, we used metagenomeSeq (Paulson et al., 2013), after controlling for (including as variables) vegetation community (categorical variable), pH (continuous variable), and %C (continuous variable), resulting in an estimate of the log_2_-fold change in the abundance of each OTU in burned *vs*. unburned plots, across samples.

We tested whether there were differences in estimated richness between years and burn severities using breakaway (Willis and Bunge, 2015), using the breakaway_nof1 function, which does not require that singletons be included. The algorithm applies different statistical models depending on the limitations of the data structure for each sample, so we only compared samples to each other where the same statistical model was selected. We interpreted treatments with non-overlapping 95% confidence intervals as being significantly different.

In order to determine whether fungal community and understory vegetation community compositions were correlated, we used a Mantel test on Bray-Curtis dissimilarities (as implemented in vegan (Oksanen et al., 2021), and using the Spearman’s rank correlation) of understory vegetation community composition data for the same sites (as published in Dawe et al., 2022) and relative abundances of fungal OTUs, considering organic and mineral horizons separately.

## 3. Results

### 3.1 Community-level patterns persist one vs. five years post-fire

Five years post-fire, vegetation community, moisture regime, pH, total C, texture (measured one year post-fire), and burned/unburned all remained significant predictors of fungal community composition, with similar R^2^ values (Table 1). The R^2^ for burned/unburned decreased between one and five years post-fire, while R^2^ for vegetation community increased. In a model with both years of data, years since fire (categorical variable) had an R^2^ of 0.05 (p=0.001), of a similar magnitude to burned/unburned in the combined years model (R^2^ = 0.05, p=0.001).

**Table 1.** PERMANOVA results one year (2015) and five years (2019) post-fire. Parameters are listed in order of inclusion in model.

| Parameter | R<sup>2</sup> |  | p-value |  |
| --- | --- | --- | --- | --- |
|  | 2015 | 2019 | 2015 | 2019 |
| Vegetation Community | 0.09 | 0.10 | 0.001 | 0.001 |
| Moisture Regime | 0.04 | 0.03 | 0.001 | 0.001 |
| pH | 0.06 | 0.05 | 0.001 | 0.001 |
| Total C (%) | 0.04 | 0.04 | 0.001 | 0.001 |
| Sand (%) (in 2015) | 0.02 | 0.02 | 0.001 | 0.001 |
| Burned / Unburned | 0.08 | 0.05 | 0.001 | 0.001 |
| Residuals | 0.68 | 0.71 |  |  |

For upland burned sites where vegetation transitions had occurred, five year post-fire dominant overstory vegetation was a stronger predictor of fungal community composition than pre-fire overstory vegetation (PERMANOVA, R^2^ of 0.06 *vs*. 0.04, p = 0.004 and p = 0.02, respectively).

There were not significant differences in estimated richness across burn severities for either year (Supplemental Figure 1). Samples collected five years post-fire tended to have higher estimated richness, including in the unburned samples.

Five years post-fire, fungal communities in burned sites were not significantly more similar to those of unburned sites than they were one year post-fire (Fig. 2; Mann-Whitney U test, p = 0.47).

**Figure 2.**
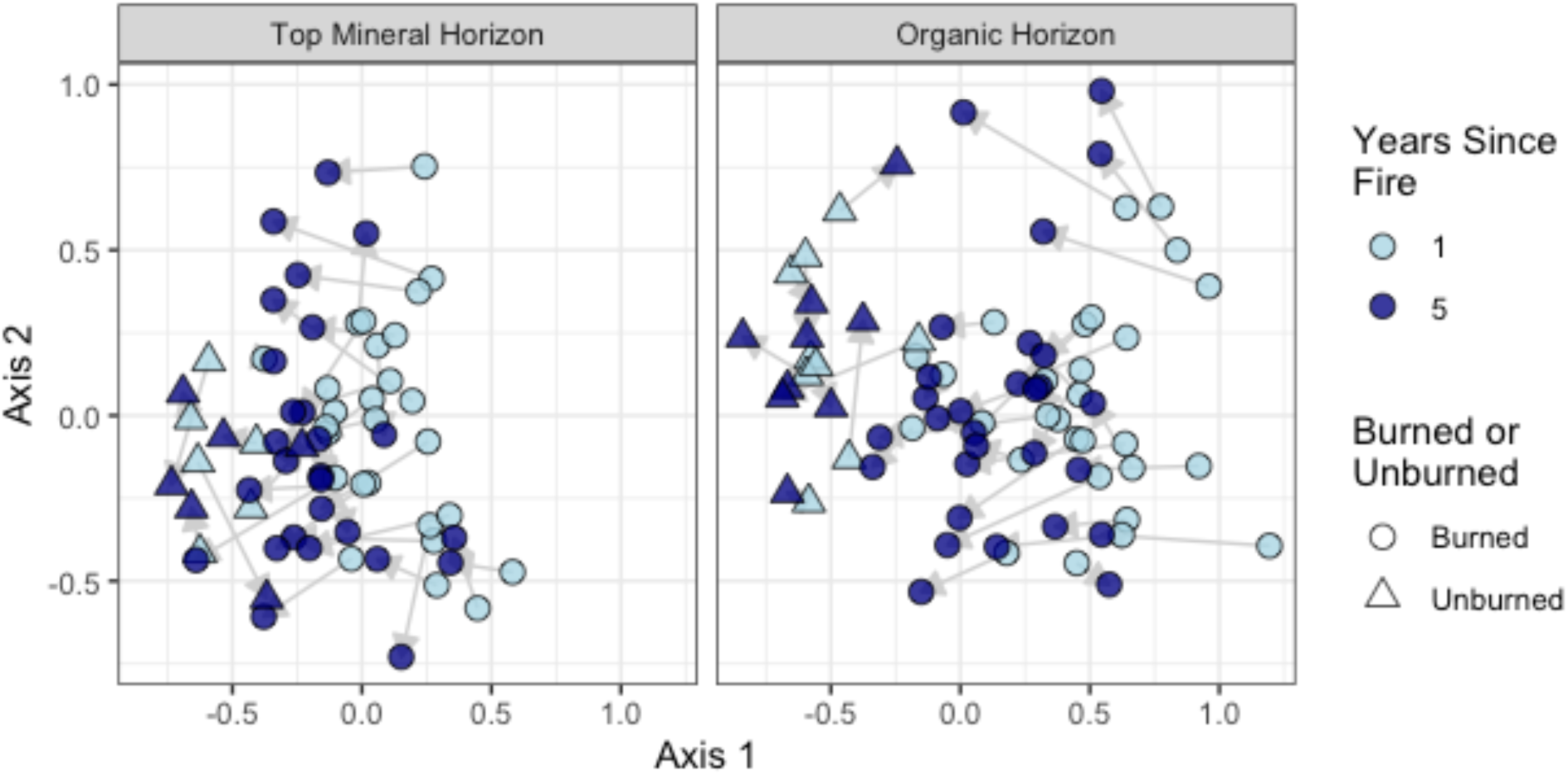
First two axes of NMDS of Bray-Curtis dissimilarities of Hellinger-transformed OTU table, separated by organic vs. mineral horizons (k=3, stress = 0.17). Triangles represent unburned sites, circles represent burned sites, and grey arrows join the same site between years 1 (light blue) and 5 (dark blue) post-fire.

Burn severity was not significantly correlated with the degree to which fungal communities changed between one and five years post-fire (Supplemental Figure 2, Kruskal-Wallis test (burned samples only), p=0.07)

In organic horizons, fungal community composition was not significantly correlated with understory vegetation community composition one year post-fire (simulated p value=0.05), but fungi and understory vegetation communities were relatively strongly correlated five years post-fire (Mantel’s R=0.51, simulated p value=0.001). In mineral horizons, there was a significant correlation with understory vegetation both one year (Mantel’s R =0.28, simulated p value=0.002) and five years (Mantel’s R =0.18, simulated p value=0.04) post-fire.

### 3.2 Similar taxa remain enriched five years post-fire, with variation in specific OTUs

The ratio of *Ascomycetes* to *Basidiomycetes* was significantly and positively correlated with burn severity index (p=0.001) and decreased between one and five years post-fire (p<0.001). However, it is critical to note that this ratio also decreased in unburned plots (Figure 3).

**Figure 3.**
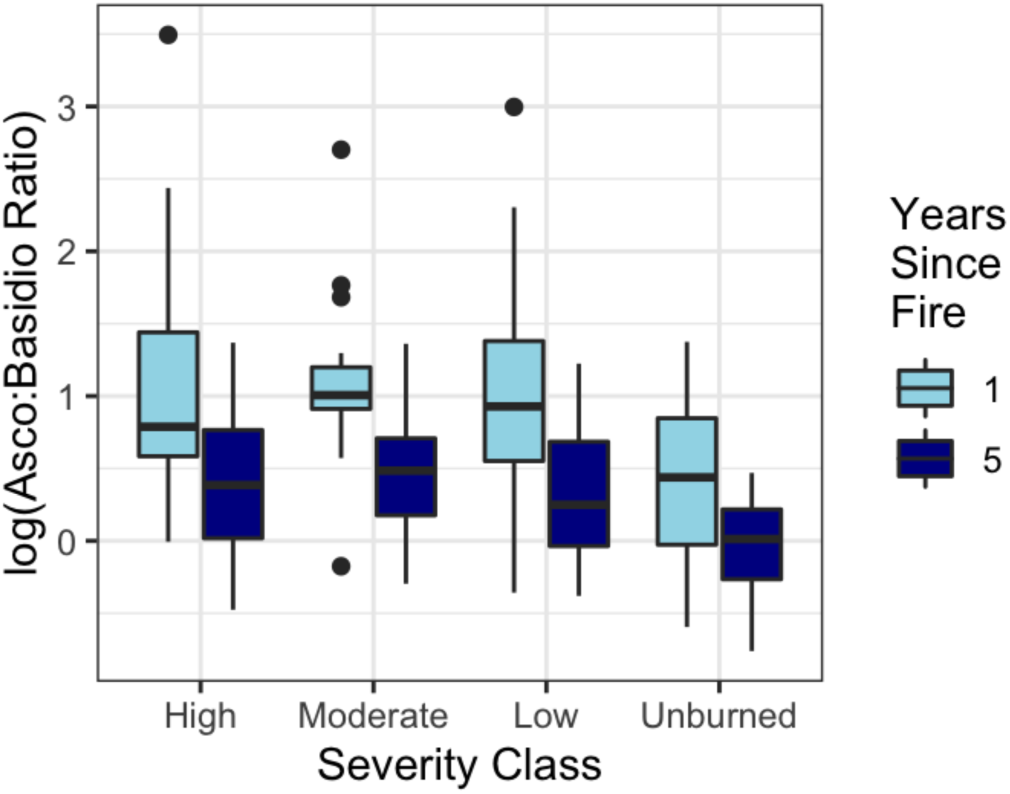
Log-transformed ratio of relative abundance of *Ascomycetes* to *Basidiomycetes* one (light blue) and five (dark blue) years post-fire across burn severities.

Five years post-fire, we found more enriched (72 *vs*. 52 one year post-fire) and more depleted taxa (59 *vs*. 39 one year post-fire) in burned *vs*. unburned sites. However, there were also more taxa overall observed five years post-fire (3607 *vs*. 1497 one year post-fire; Supplemental Figure 1), meaning a smaller proportion of total observed OTUs were enriched or depleted in the five year dataset. Responsive taxa generally had similar responses for both years, with 28 taxa being significantly enriched with fire both years, and 20 taxa being significantly depleted both years (Figure 4; Supplemental Table S2).

**Figure 4.**
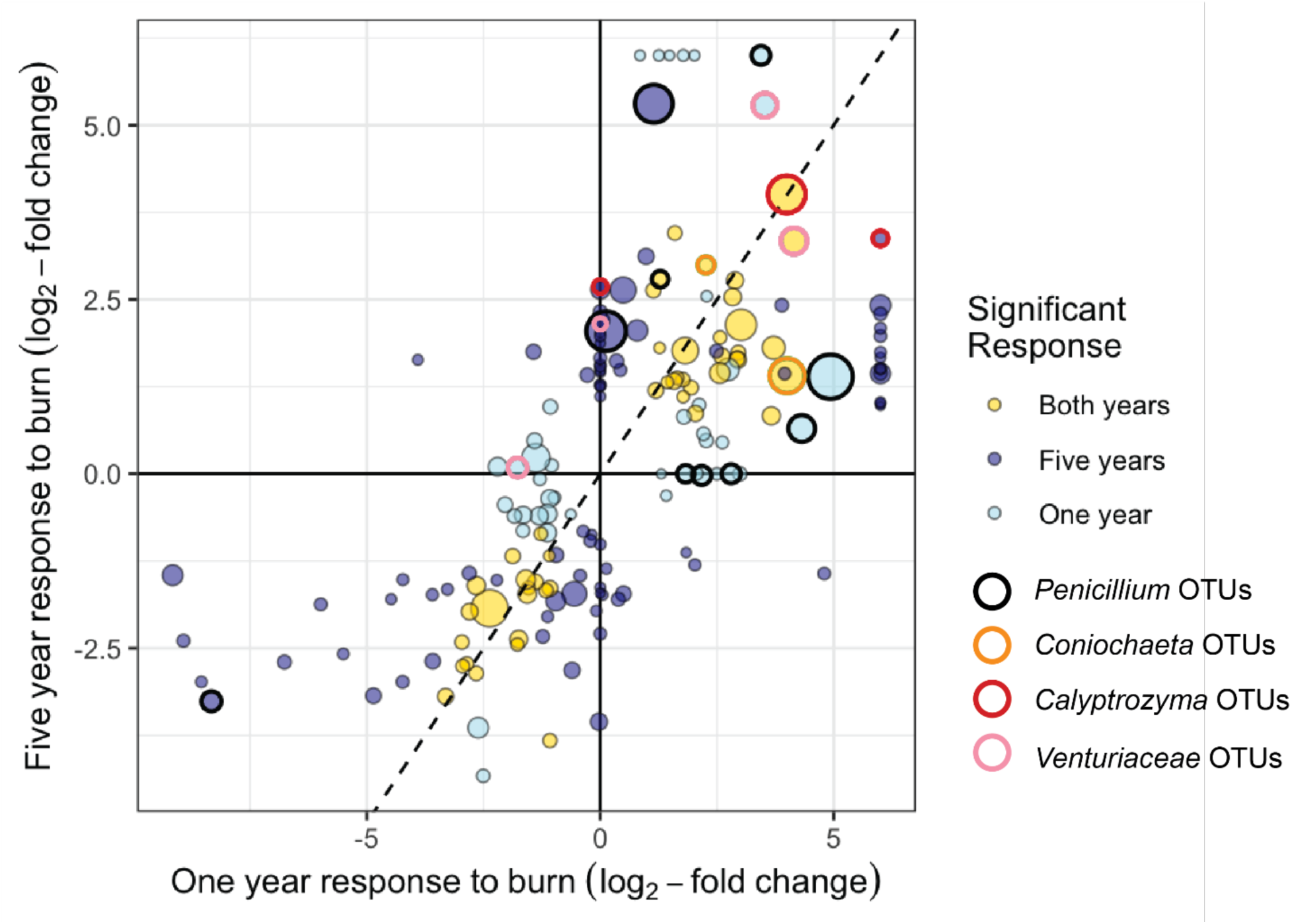
Degree of enrichment in burned vs. unburned sites, five years *vs*. one year post-fire for all responders. Each point indicates a single OTU, and colours indicate the years in which their enrichment (or depletion) in burned samples was statistically significant. Points are generally sized by their mean relative abundances across datasets. Taxa that were significant in one year, but not detected in burned, unburned, or both treatments in the other year, could not be tested for significance. In order to include them in the plot, they were given arbitrary values (+6 if found in burned plots but never detected in unburned plots; 0 if not detected in any plots; −6 if they were found in burned but never detected in unburned plots for that year). Outlined points highlight OTUs from the genera *Penicillium*, *Coniochaeta*, *Calyptrozyma*, and the family *Venturiaceae* (which includes the genera *Fusicladium*/*Tyrannosorus*/*Venturia*).

There were seven *Penicillium* taxa that were enriched one year post-fire, and three that were enriched five years post-fire, but only one of these was shared between the two years. These *Penicillium* OTUs that were responsive only at five years post-fire were less than 97% similar to all of the one-year taxa, suggesting that they are truly distinct OTUs. A selection of abundant *Penicillium* OTUs is plotted in Figure 5, which illustrates different types of responses to fire in these OTUs from the same genus.

**Figure 5.**
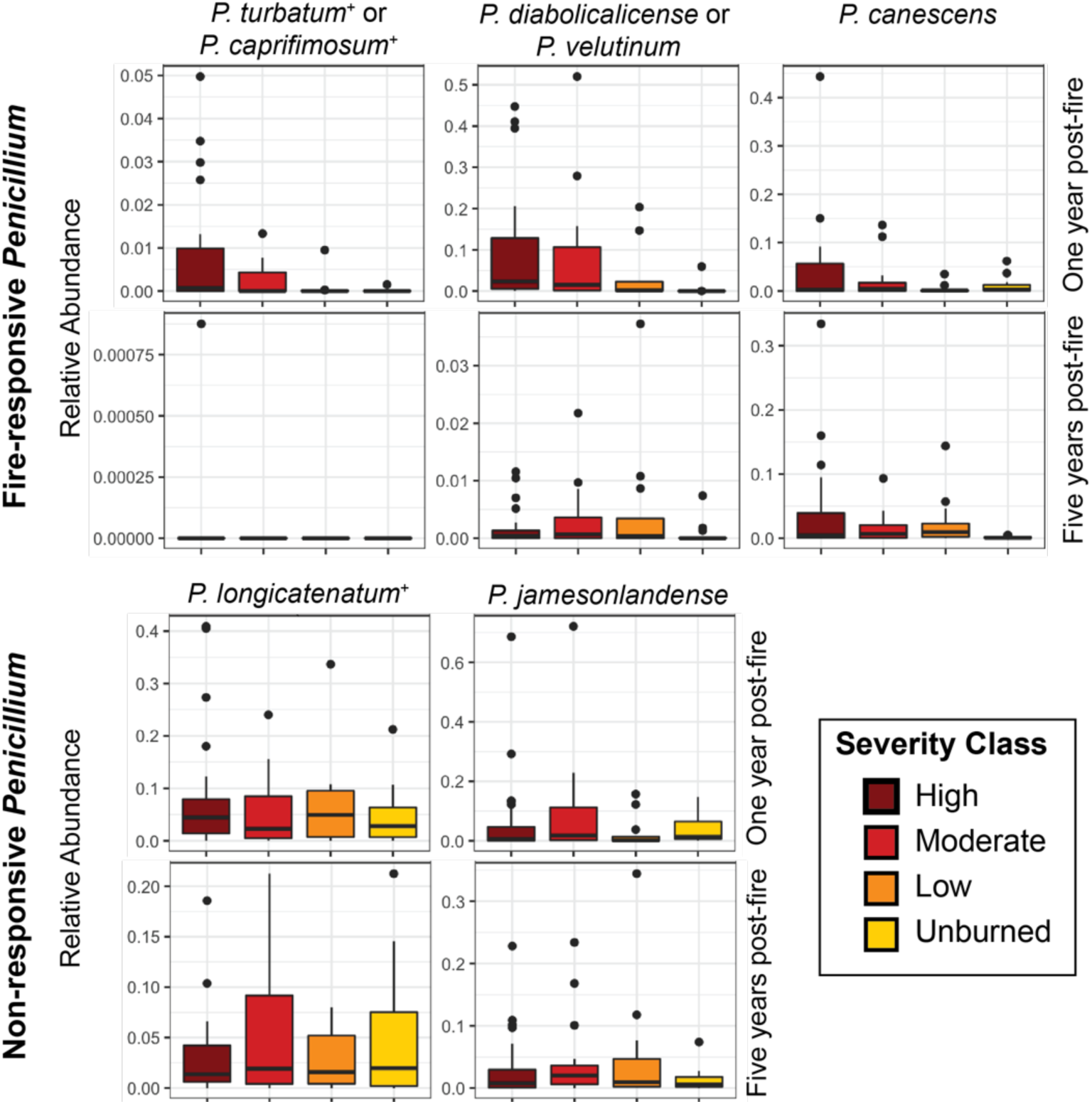
Relative abundances of selected *Penicillium* OTUs one (upper row) and five (lower row) years post-fire across all samples. These selected *Penicillium* OTUs include three taxa that were fire-enriched in at least one year (top) as well as two generally abundant but not fire-responsive OTUs (bottom). OTUs are labelled with their UNITE-matched taxonomy, or best NCBI BLAST matches. + indicates OTUs that are at least a 99%ID match for *Penicillium* isolates described in Day et al. (2020). Note different scales on y-axes.

### 3.3 Enrichment of saprotrophs in burned soils decreases five years post-fire, while depletion of symbiotrophs intensifies

“Highly probable” saprotrophs were more abundant at burned than unburned sites one year post-fire in both mineral and organic horizons, but this effect was no longer significant five years post-fire (Figure 6A). There were no significant differences in the relative abundances of “highly probable” pathotrophs at burned *vs*. unburned sites (Figure 6B). “Highly probable” symbiotrophs emerged as being less abundant at burned than unburned sites in both mineral and organic horizons five years post-fire (Figure 6C). The saprotroph trends are largely driven by OTUs from the genus *Penicillium* (classified as a “Dung Saprotroph-Undefined Saprotroph-Wood Saprotroph”), whereas the trends in symbiotrophs are largely driven by ectomycorrhizal fungi (particularly from the genera *Cortinarius*, *Piloderma*, and *Inocybe*) and endophytes (particularly dark septate fungus *Leptodontidium* in the O horizons).

**Figure 6.**
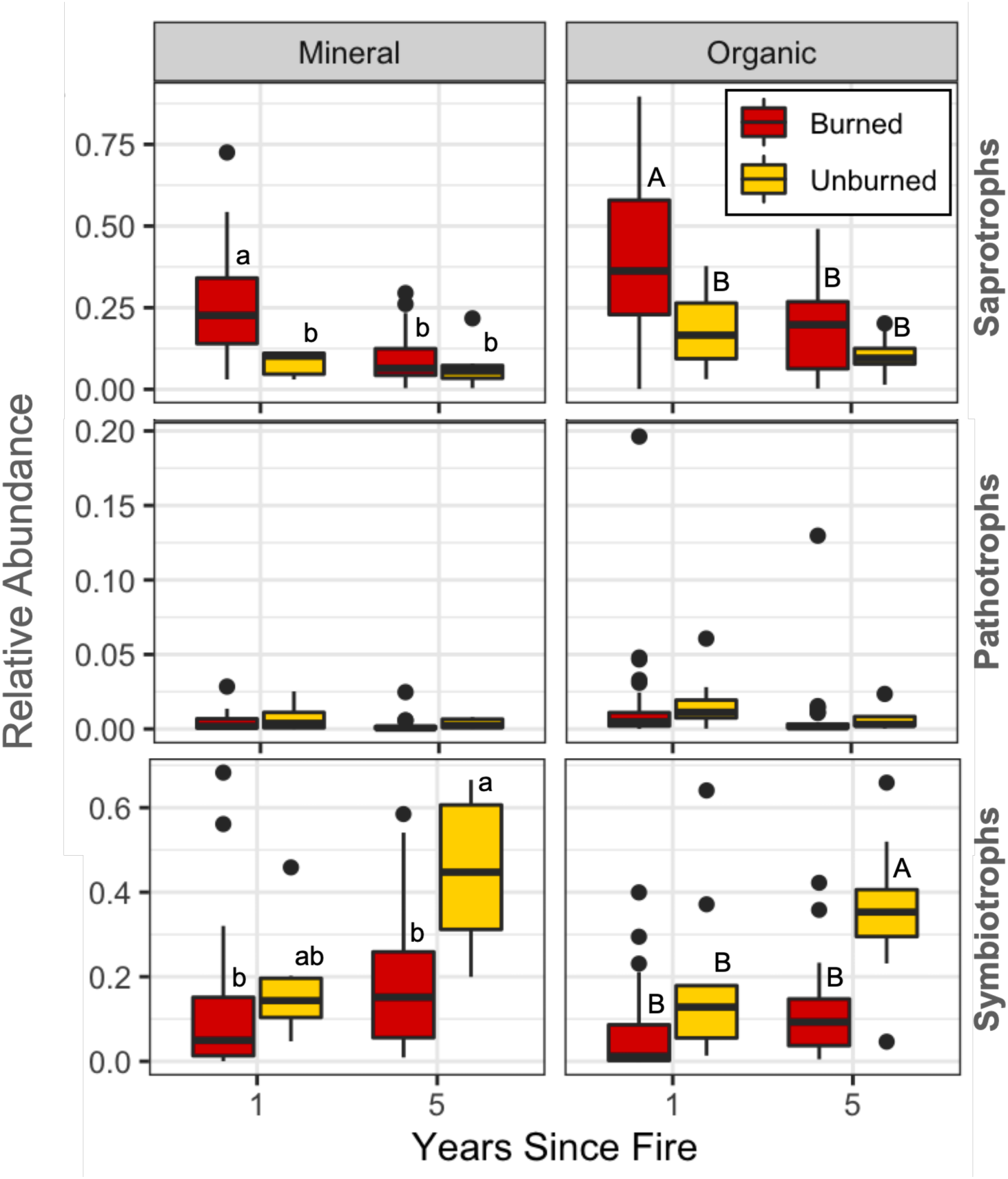
Relative abundance of “highly probable” saprotrophs, pathotrophs, and symbiotrophs in burned (red) and unburned (yellow) samples, one and five years post-fire, in mineral (left panels) and organic horizons (right panels). Letters indicate significant differences within a given panel (ANOVA, Tukey’s HSD).

## 4. Discussion

### 4.1 The influence of burning on fungal community composition declines five years post-fire

Consistent with our hypothesis, similar factors, including vegetation community, moisture regime, pH, percent total C, and percent sand, continued to structure fungal community composition five years post-fire (Table 1). This is broadly consistent with our findings in the bacterial/archaeal communities of the same samples (Whitman et al., 2022). The waning influence over time of whether a site was burned or not seems to suggest that the direct effects of burning are also decreasing, or that the effects of the fire are diverging over time, from one site to the next, so that simple factor is not as predictive as it was one year post-fire. This would be consistent with our findings from a paired laboratory-field study of bacterial/archaeal communities, where we found that fast growth was an important trait one year post-fire, but its importance (*i.e.*, the proportion of the community made up of fast growers) almost completely disappeared by five years post-fire (Johnson et al., 2023). After a fire, in the upper soil horizons that reach sufficient temperatures or heat exposure to kill large portions of the microbial community (Pressler et al., 2018; Pingree and Kobziar, 2019), a niche space is opened, paving the way for secondary succession. Because temperatures attenuate rapidly with soil depth during a fire (Bruns et al., 2020), a robust source of potential “inoculum” for the depopulated upper horizons persists below the surface, while aerial dispersal offers colonization potential from above (Kobziar et al., 2018; Walters et al., 2022). Thus, one major effect structuring the post-fire community in these ecosystems – the short-term dominance of fast-growing organisms – seems to disappear within years and may help explain the waning influence of whether a site is burned on fungal community composition.

A second potential explanation for the decreasing influence of fire on fungal community composition is that fungal communities at different sites may initially converge in composition to some extent post-fire, but then experience divergent paths post-fire. For example, the burn may initially select for a smaller subset of fire-adapted taxa, such as taxa that are resistant to or even stimulated by heat (Nguyen et al., 2013). As the broader ecosystem recovers post-fire, different sites may follow different trajectories, depending on factors such as the previous vegetation and soil conditions, and the specific burn conditions and other local factors. For example, the O horizon can be nearly completely combusted during a burn (E. Whitman et al., 2019), and as organic matter re-accumulates, the influence of the local vegetation community from which the litter is derived (*e.g.*, its chemistry, rate of formation, or physical properties) may begin to exert more influence on the fungal community (Hart et al., 2005). This is consistent with the observation that the fraction of the fungal reads representing putative saprotrophs (“Highly Probable”) was significantly higher in burned than unburned sites one year post-fire but decreased to match that of the unburned sites five years post-fire, in both organic and mineral horizons (Figure 5). These taxa may be capitalizing on the fire-killed organic matter inputs immediately post-fire, while this effect declines significantly four years later. It is also consistent with our finding that understory vegetation communities were significantly correlated with fungal communities in the O horizon five years post-fire but were not significantly related one year post-fire. This mechanism for waning influence of whether a site was burned or not would be intensified when there are divergent vegetation communities. Fire-mediated overstory vegetation transitions, often from conifer to aspen-dominated, have been documented at many of these sites (Dawe et al., 2022). If these transitions alter the fungal community, via shifts in mycorrhizal fungi associations with the new dominant vegetation or fungi that thrive on different litter chemistries, such changes would be expected to emerge over a few years, as the vegetation re-establishes. This was supported by the finding that – for sites where vegetation transitions occurred – the post-fire overstory vegetation was a better predictor for fungal community composition than pre-fire overstory vegetation.

We did not formulate specific hypotheses about richness, partly because we believe that microbial richness is only rarely a constraint on most functions of interest. Additionally, we expect effects on richness might be relatively transient, and highly dependent on sampling depth and fire severity. However, we did test for differences in richness and found a lack of significant differences with burn severity (Supplemental Figure 1) five years post-fire, which was consistent with our findings one year post-fire. Samples from the five years post-fire sites had higher richness estimates, but because increases in estimated richness were observed in the unburned samples as well as burned (Supplemental Figure 1), we do not interpret the increase in estimated richness as representing recovery post-fire, but, rather, as likely indications of seasonal fluctuations (Taylor et al., 2010; Averill et al., 2019). It could also reflect the greater sequencing depth five years post-fire (21.9k vs. 62.6k mean reads per sample, 1 and 5 years post-fire, respectively), but the statistical algorithms used to estimate richness should help account for such variation (Willis and Bunge, 2015).

### 4.2 Resilience in fungal community composition not evident after five years

Whereas burned bacterial/archaeal communities became more similar to unburned sites between one and five years post-fire, regardless of burn severity (Whitman et al., 2022), we did not find evidence for resilience in the fungal communities, counter to our hypothesis (Figure 2). This lack of resilience was observed across burn severities, and is even observed at the level of individual taxa: again, in contrast to bacteria/archaea, no OTUs that were significantly depleted one year post-fire recovered to the point of being significantly enriched five years post-fire (Figure 4). While the *Ascomycetes* to *Basidiomycetes* ratio – a metric roughly related to increased stress – did decrease between one and five years post-fire (Figure 3), this effect was also observed in the unburned sites, and thus more likely represents inter-annual variation in environmental conditions than recovery. Specifically, the one year timepoint from 2015 may still be reflecting the very dry conditions of 2014, the year of the fires (Dawe et al., 2022), given that drought stress is associated with higher *Ascomycetes* to *Basidiomycetes* ratios (Wan et al., 2023).

Together, this could indicate that fungal communities are less resilient than bacteria/archaea after a burn, taking longer to recover. If this is the case, it suggests that fungi may be more closely tied to soil properties or vegetation communities, which can take years to decades to recover after a burn. For example, one might predict that fire-killed organic matter would provide an important resource for fungi for many years, potentially maintaining an enrichment of saprotrophs. However, we do not observe this phenomenon, with saprotroph enrichment in burned sites disappearing by five years post-fire (Figure 5A). Furthermore, the relative predictive power of factors that structure microbial communities between one and five years post-fire generally changed in similar ways for bacteria/archaea (Whitman et al., 2022) as for fungi (Table 1), not offering clear support to a specific factor that might be maintaining this difference in apparent resilience. Notably, the fungal and bacterial/archaeal data were generated from the same DNA extractions, ruling out a number of possible methodological artifacts as explanations.

It is also notable that we are observing these responses to wildfire against a backdrop of long-term climatic changes in the region. Between 1965 and 2020, much of the study region experienced significant increases in summer maximum temperatures of a mean of 1.1°C and an overall decrease in summer vapour pressure deficit, reflecting increasingly dry conditions (Dawe et al., 2022). If changes in climate interact with post-fire recovery of fungal community composition, that could affect perceived resilience. Additionally, if the DNA of dead fungi, or “relic DNA” (Carini et al., 2016) is more persistent than that of bacteria and archaea, then its persistence could obscure trends toward recovery in the living communities. Despite changes in climate, the region still experiences long cold winters, which should slow apparent changes in total DNA.

There may also be influential factors that were not measured in this study – in particular, soil nutrients such as available nitrogen or phosphorus – that could potentially affect fungal recovery rates. Because mineral nitrogen concentrations and fluxes in a soil can be highly variable over time (and space), a single timepoint measurement from our soil samples might not be particularly informative. However, nitrogen stocks and fluxes have often been observed to be affected by fire (Pellegrini et al., 2017), and fungal communities may respond differently to these changes than bacterial communities (Smithwick et al., 2012). For example, at the extremely coarse scale of a global dataset, Bahram et al. (2018) reported that C:N was significantly positively correlated with fungal abundance relative to bacteria, but also positively correlated to both bacterial and fungal biomass overall, and that bacterial to fungal ratios were positively associated with nitrate. Fires can affect soil available P (Souza-Alonso et al., 2024), with effects potentially persisting over years to decadal scales (Zhou et al., 2025), which could then alter relationships between plants and mycorrhizal fungi. Still, an explanation relying on effects on available nitrogen or phosphorus may not be strong, given that we would expect mineral nutrients to vary with burn severity (e.g., (Turner et al., 2007)), but we found a lack of resilience across all levels of burn severity.

Further mechanistic work might help illuminate the reasons for lack of significant recovery in fungal community composition five years post-fire. Long-term monitoring will allow us to see whether this effect continues to persist throughout the decade post-fire. For example, despite the lack of a clear resilience, we did not find any relationship between the degree to which fungal community composition changed and burn severity (Supplemental Figure 2). This could suggest that severely burned sites are not necessarily “impaired” in some consistent way, compared to sites with low-severity burns. Additionally, the general – non-statistically significant – trend for changes in fungal community composition in burned sites does seem to be toward increasing similarity to unburned sites (Figure 2).

### 4.3 Specific fire-responsive taxa have different patterns of enrichment

Fungal taxa enriched in burned samples included taxa from our predicted genera, with *Penicillium*, *Calyptrozyma*, and *Coniochaeta* OTUs remaining significantly enriched five years post-fire. We did not observe an enriched *Fusicladium* between both years, but did detect an OTU from the family *Venturiaceae*, which was a 100% ID match to *Tyrannosorus hystrioides* type material (identified as *Venturia hystroides* (Crous et al., 2007)). *Venturia* and *Fusicladium* represent teleomorphs and anamorphs of a monophyletic group (Beck et al., 2005). Most of the persistent responders were identified as microfungi, yeasts, or facultative yeasts, with the two exceptions being a “possible” ectomycorrhizal agaricoid *Plicaria* and a “probable” pezizoid white rot fungus *Pholiota* (Supplemental Table S2). Many of these fire-enriched taxa are likely playing saprotrophic roles (*e.g.*, *Penicillium*), with some potential pathotrophs (*e.g.*, fungal parasite *Cystobasidium*), and some having “possible” or “probable” roles as some combination of saprotrophs, pathotrophs, and/or symbiotrophs (*e.g.*, *Coniochaeta*). All of these roles would be consistent with fungi capitalizing on an influx of dead organic matter post-fire (Figure 6A), and possibly taking advantage of injured plants, as pathogens. The trends in putative symbiotrophs (Figure 6C) are more difficult to explain. The changes in the unburned sites are largely driven by increases in *Cortinarius*, which is associated with more mature stands, and may reflect changes in its host plants’ performance between the two years (Fernandez et al., 2016). Regardless, it is important to note that any of the putative traits discussed in this paragraph are not sufficient alone to drive enrichment – fire-depleted taxa include OTUs with these same trait assignments.

Fox et al. (2022) outline three (overlapping) categories of pyrophilous fungi: fire-resistant (“heat resistant for some aspect of their life cycle”), fire-responsive (“triggered to grow and/or fruit by heat or by chemical changes caused by heat”), and fire-adapted (“with specific traits related to fire and that may require fire to complete their life cycle”). Because the two axes in Figure 4 represent enrichment or depletion shortly after the fire on the x-axis, and later enrichment or depletion on the y-axis, the space doesn’t map perfectly on to Fox et al.’s categories. “Fire-resistant” and “fire-adapted” taxa may or may not become enriched, especially since the axes in Figure 4 are related to changes in relative, not absolute, abundance. For example, if fungi are fire-resistant but not fire-responsive, then they would generally not be identified as being enriched (or only be slightly enriched one year post-fire). However, many “fire-responsive” taxa should be enriched one and/or five years post-fire. This way of plotting fungal OTUs could allow us to identify three categories of fire-responsive taxa: emergently fire-responsive, persistently fire-responsive, and fleetingly fire-responsive (yellow, orange, and red areas in Figure 7). Similarly, for depleted taxa, we might classify them as fleetingly fire-sensitive, persistently fire-sensitive, and emergently fire-sensitive (dark to light blue areas, Figure 7).

**Figure 7.**
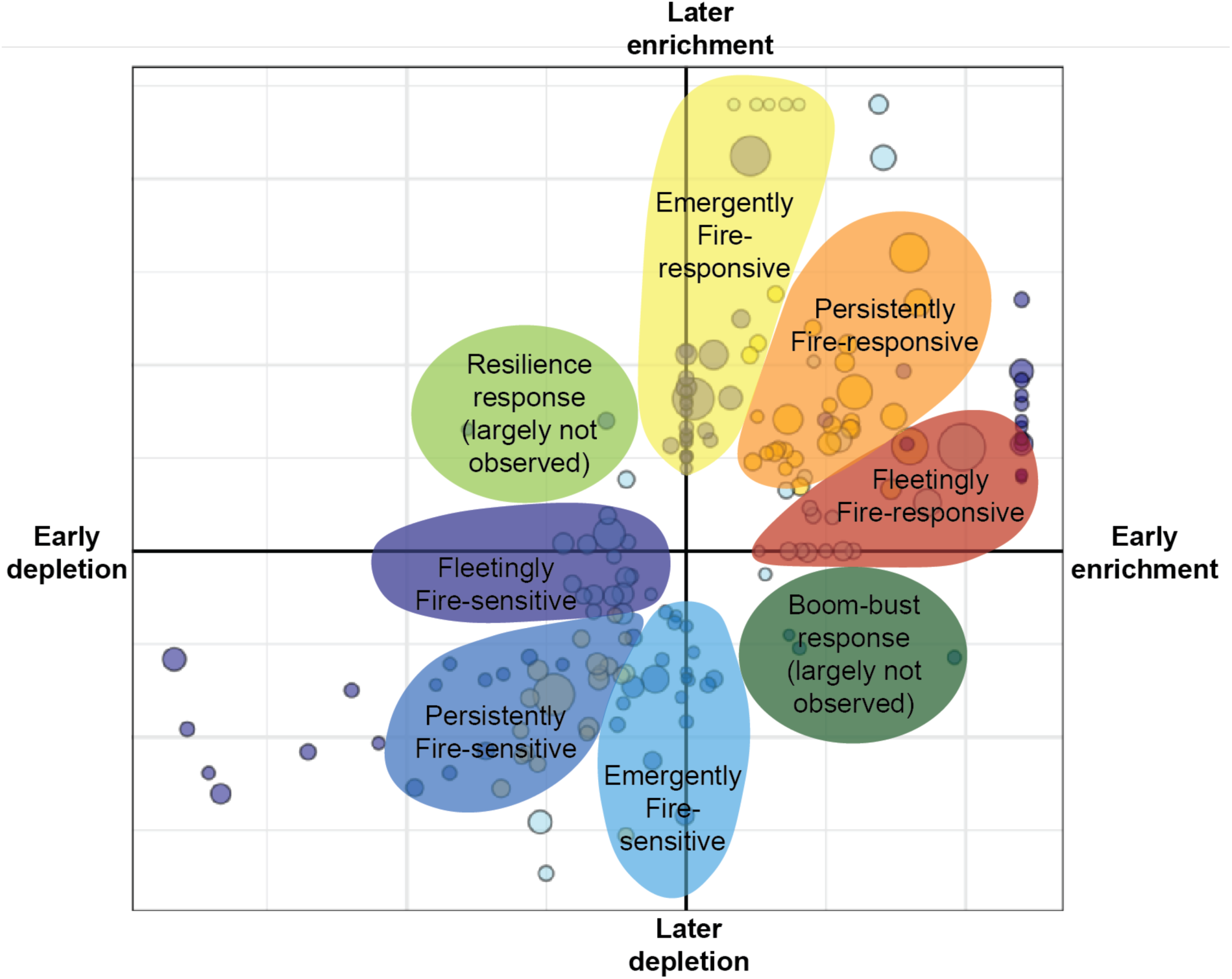
Conceptual illustration of putative fire-response traits mapped on to axes of Figure 4.

These different sub-categories might indicate different underlying reasons for enrichment or depletion. For example, the fleetingly fire-responsive taxa (red, Figure 7), rapidly increase in relative abundance one year post-fire, but do not remain enriched. These taxa would be consistent with fast-growing copiotrophs taking advantage of a post-fire pulse in nutrients and may be equivalent to the “fast-growing” bacteria that we identify one year post-fire (Johnson et al., 2023). Conversely, the emergently fire-responsive taxa (yellow, Figure 7) only emerge as being enriched five years after the fire. These taxa may be responding to the post-fire environment – e.g., thrive under the new soil chemical or physical conditions – but are not particularly fast-growing. Together, the emergently and persistently fire-responsive tax (yellow and orange, Figure 7) would include some of the fungal analogues for the vegetation species that dominate when vegetation transition occurs after fire (Dawe et al., 2023), although our method used here would not identify organisms that are completely absent in the unburned sites.

The fleetingly fire-sensitive taxa (dark blue, Figure 7) may reflect taxa that are directly killed by the fire, whereas the emergently fire-sensitive taxa (light blue, Figure 7) may reflect taxa that survived the fire, but did not fare well under post-fire conditions, and thus became depleted over a longer period. Interestingly, over the span of this experiment, we observed almost no individual taxa that would be identified as being resilient (i.e., depleted one year post-fire, but enriched at five years post-fire; light green area, Figure 7). As noted above, this is in contrast with our findings for bacteria (Whitman et al., 2022), and may reflect the faster timescales over which some bacteria recover post-fire. We also observe very few taxa in the opposite quadrant (“boom-bust response), where they are initially enriched but subsequently become depleted (dark green area, Figure 7). For all categories, it will be valuable to assess their consistency over longer timescales. Future sampling at these sites, a decade or more post-fire, will enable evaluation of whether five-year post-fire responses align with those observed at ten years or longer.

While 28 taxa remained enriched both one and five years post-fire, some of the strongest responders for each year changed. When we first recognized that some of the strongest and most abundant responders were all classified as *Penicillium*, we initially questioned whether we had inadvertently split different ITS2 copies within the same genome into different taxa. However, alignment and taxonomy suggest that there are indeed distinct *Penicillium* OTUs, which exhibit different patterns of enrichment (Figure 5). For example, the putative *Penicillium diabolicalicense* makes up large fractions of the sequenced reads one year post-fire, but strongly falls back five years post-fire, by a factor of roughly ten. The putative *Penicillium turbatum* mirrors this pattern of a strong, short-term response, becoming almost undetectable five years post-fire. In contrast, the putative *Penicillium canescens* remains similarly enriched in burned *vs*. unburned sites both one and five years post-fire. Meanwhile, other *Penicillium* spp. represent persistently large fractions of communities across burned and unburned sites (Figure 5).

These patterns highlight simultaneously the need to apply fine-resolution techniques that allow for the detection of closely-related different species or strains that have distinct ecological patterns, but also the need to consider consistent responses at coarser taxonomic/genetic resolution. In this dataset, numerous *Penicillium* spp. are identified as positive fire responders, but different species/strains have different response patterns across different sites and time since fire. Future research could focus on disentangling these responses to identify the specific environmental factors or traits driving these distinct patterns.

### 4.4 Summary and Future Directions

Returning to the same sites one and five years post-fire, we found that although the importance of whether a site was burned in structuring soil fungal communities decreased five years post-fire, fungal communities failed to show resilience in community composition, possibly due to strong interactions with living overstory and understory vegetation as well as fire-killed plant matter. Assessing the multi-year response of individual taxa to fires underscores that neither genus-level taxonomy nor putative trophic assignment is sufficient to consistently explain post-fire response. Future manipulative and observational studies will help us continue to dissect the multiple factors and traits structuring fungal responses to fires.

## Data Availability

Sequencing data are deposited in the NCBI SRA under PRJNA564811 (public) and PRJNA1243682 (to be made public upon publication). Code associated with processing sequencing data, analysis, and figures for this paper can be found at https://github.com/TheaWhitman/WoodBuffalo1yr5yr_fungi.

## Acknowledgements

Thanks to Marc-André Parisien, Mike D. Flannigan, Daniel K. Thompson, and Élyse Mathieu for their contributions in the field and/or to the original study design. This work was supported by the Government of the Northwest Territories, which provided in-kind and financial support for the field campaign that produced these data; Parks Canada Agency and Jean Morin at Wood Buffalo National Park provided in-kind support during fieldwork; the U.S. Department of Energy helped support T.W. and J.W. during data collection [DE-SC0016365; DE-SC0020351] and the U.S. National Science Foundation helped support T.W. during data analysis [2045864].

## Supplementary Data

Supplementary data are associated with this article.

