## Supplemental Information for "Resilience not yet apparent in soil fungal communities of the boreal forest from one to five years after wildfire across a severity gradient"

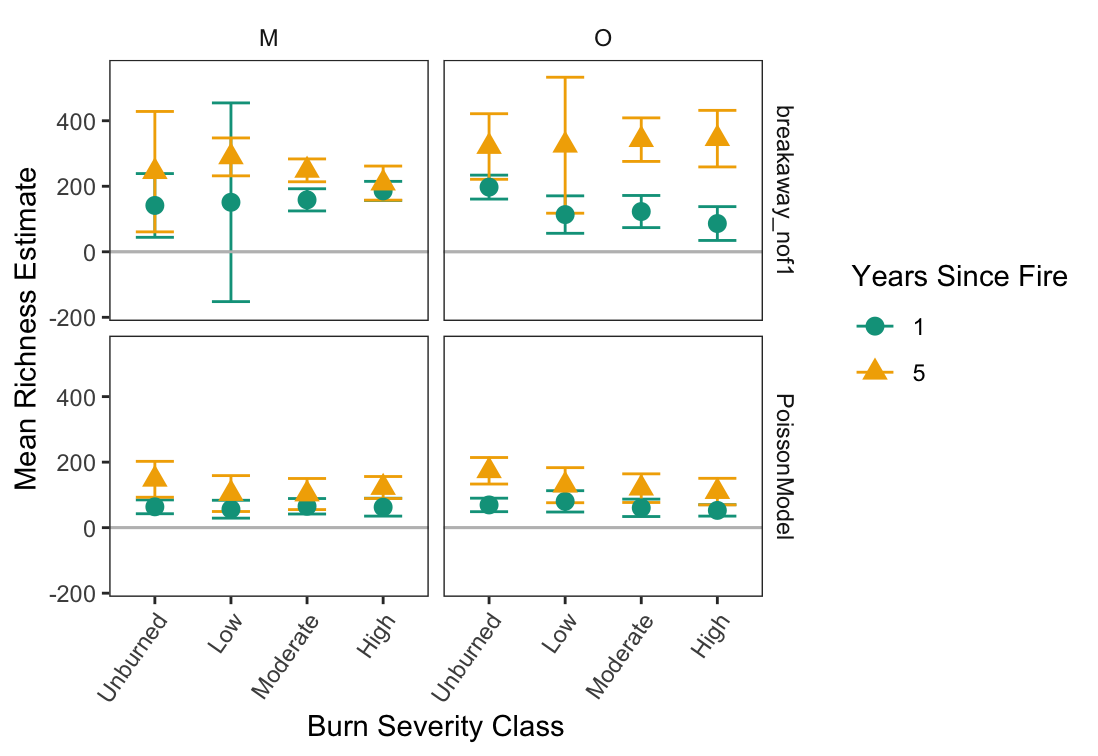

Supplemental Figure 1. Estimated richness one and five years post-fire across burn severity index categories. Error bars represent 95% confidence intervals (±1.96 standard error), where non-overlapping bars suggest significant differences. The panels separate data for mineral (“M”) and organic (“O”) soil horizons and the two different statistical models used (different samples’ data structure required different statistical models).

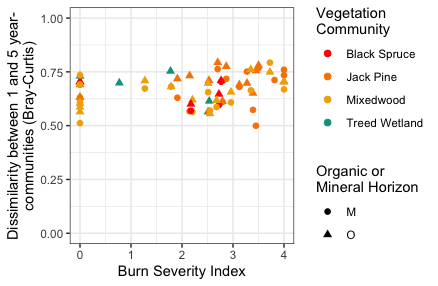

Supplemental Figure 2. Bray-Curtis dissimilarity on Hellinger-transformed relative abundances between paired sites one *vs*. five years post-fire vs. burn severity index.

| Supplemental Table 1. Primers used in this study to amplify ITS2 region (Taylor et al., 2016) using Illumina MiSeq 2x300 Paired Ends sequencing (Kozich et al., 2013) | |
| --- | --- |
| **Primer** | **Sequence [**Illumina adaptor *Barcode* **Pad and linker** Primer] |
| ITS4 | AATGATACGGCGACCACCGAGATCTACAC*XXXXXXXX***TATGGTAATTAA**AGCCTCCGCTTATTGATATGCTTAART |
| 5.8S | CAAGCAGAAGACGGCATACGAGAT*XXXXXXXX***AGTCAGTCAGGG**AACTTTYRRCAAYGGATCWCT |
| Read 1 seq | TATGGTAATTAAAGCCTCCGCTTATTGATATGCTTAART |
| Read 2 seq | AGTCAGTCAGGGAACTTTYRRCAAYGGATCWCT |
| Barcode seq | AGWGATCCRTTGYYRAAAGTTCCCTGACTGACT |

Supplemental Table 2. Strong and relatively abundant fire-responders.csv
